# Modulation of the Gut Microbiome by Novel Synthetic Glycans for the Production of Propionate and the Reduction of Cardiometabolic Risk Factors

**DOI:** 10.1101/2022.04.04.487010

**Authors:** Yves A. Millet, Jeffrey Meisner, Jie Tan, Adarsh Jose, Eric Humphries, Kelsey J. Miller, Charlie Bayne, Megan McComb, Michael Giuggio, Camille M. Konopnicki, David B. Belanger, Lingyao Li, Han Yuan, Madeline Rosini, Hoa Luong, Jared Martin, Zhengzheng Pan, C. Ronald Kahn, Johan E.T. van Hylckama Vlieg

**Affiliations:** Kaleido Biosciences, 65 Hayden Ave, Lexington, MA 02421, United States; Integrative Physiology and Metabolism, Joslin Diabetes Center, Harvard Medical School, Boston, MA, 02215, United States

## Abstract

**Background:** Increasing evidence indicates that an altered gut microbiome participates in the development of cardiometabolic syndrome and associated risk factors, such as insulin resistance, dyslipidemia, and obesity, and that targeting the gut microbiome is a promising strategy to lower the risk for cardiometabolic diseases. Part of this reduction is mediated by specific metabolites generated by the gut microbiome. Propionate, a short-chain fatty acid (SCFA) produced from dietary glycans by certain gut microbes is known to exert multiple beneficial metabolic effects. Here, we identify KB39, a novel gut microbiome-targeting synthetic glycan selected for its strong propionate-producing capacity, and demonstrate its effects *in vivo* to reduce cardiometabolic disease using western diet-fed LDL receptor knock-out mice.

**Methods:** *Ex vivo* fermentation screening of a large library of synthetic glycan ensembles was performed using gut microbiome communities from healthy subjects and overweight patients with type 2 diabetes. A synthetic glycan identified for its high propionate-producing capacity (KB39) was then tested *in vivo* for effects on systemic, blood and cecal metabolic parameters in *Ldlr^-/-^* mice fed a western diet.

**Results:** *Ex vivo* screening of ~600 synthetic glycans using human gut microbiota from healthy subjects and patients with type 2 diabetes identified a novel glycan (KB39) with high propionate-producing capacity that increased propionate contribution to total SCFA and propionate-producing bacterial taxa compared to negative control. In western diet-fed *Ldlr^-/-^* mice, KB39 treatment resulted in an enrichment in propiogenic bacteria and propionate biosynthetic genes *in vivo* and an increase in absolute and relative amounts of propionate in the cecum. This also resulted in significant decreases in serum total cholesterol, LDL-cholesterol, and insulin levels, as well as reduced hepatic triglycerides and cholesterol content compared to non-treated animals. Importantly, KB39 treatment significantly reduced atherosclerosis, liver steatosis and inflammation, upregulated hepatic expression of genes involved in fatty acid oxidation and downregulated transcriptional markers of inflammation, fibrosis and insulin resistance with only a mild lowering of body weight gain.

**Conclusions:** Our data show that KB39, a novel synthetic glycan supporting a high propionate-producing microbiome, can reduce cardiometabolic risk factors and disease in mice and suggest this approach could be of benefit for the prevention or treatment of cardiometabolic diseases in humans.

**CLINICAL PERSPECTIVE:** **What is new?**

- A novel synthetic glycan, KB39, was selected from a library of compounds for its high propionate-producing capacity and beneficial effects on the human gut microbiome composition
- KB39 modulates the gut microbiome for high propionate production and significantly reduces cardiometabolic risk factors and disease in a murine model of cardiometabolic diseases

**What are the clinical implications?**

- KB39, delivered orally, could be of benefit for the prevention or treatment of cardiometabolic diseases in humans
- The efficacy of KB39 in mice compared to the clinical drug fenofibrate justifies further study in humans

## INTRODUCTION

In the past 40 years, cardiometabolic diseases, comprising of cardiovascular disease (CVD), type 2 diabetes mellitus (T2DM), non-alcoholic fatty liver disease (NAFLD) and other obesity-related comorbidities, have become a major global health issue and are now the leading cause of death and disability worldwide (WHO, World Health Statistics Report 2020). Lifestyle modifications have some effect to reduce cardiometabolic risk factors (CMRFs) ^1–7^, however, this is not sufficient and most patients ultimately require pharmacological therapies ^8, 9^, often with a multitude of drugs to address the many individual components of disease risk ^10, 11^, increasing the risk of side effects, drug-drug interactions, poor adherence and medication errors ^12^. Therefore, there is a great need to develop safe and effective drugs targeting multiple CMRFs simultaneously for the prevention of cardiometabolic diseases.

Poor quality western diets, high in sugars, salt, saturated fats and low in fibers, play a major role in the development of cardiometabolic diseases ^13–15^. These diets also lead to significant changes in the bacterial composition and metabolic output of the gut microbiota ^16, 17^, and increasing evidence suggest that this alteration of the gut microbiota contributes to the aggravation of multiple CMRFs, such as low-grade inflammation, insulin resistance, atherogenic dyslipidemia and fatty liver ^18^. Therefore, the gut microbiome is emerging as a promising target in the prevention against cardiometabolic diseases ^19^. Altered gut microbiome features caused by western diets and found in patients with CMRFs include an enrichment in pro-inflammatory mucosa-damaging bacteria belonging to the *Enterobacteriaceae* family and the depletion in fiber-degrading commensals leading to a reduction in the production and absorption of short-chain fatty acids (SCFAs), a class of microbial fermentation products of dietary fibers, that may have beneficial metabolic effects ^18^. Among the different SCFAs, propionate has received the most attention in metabolism and obesity research due to its multiple effects against body weight gain, fatty acid and cholesterol synthesis, insulin resistance, adipose tissue inflammation, atherosclerosis and hypertensive cardiovascular damage in mice ^20–24^.

It is now established that glycan utilization by the gut microbes is a major determinant of the composition and metabolic output of the gut microbiome ^25^ and the use of natural or enzymatically synthesized fibers can be used to modulate the gut microbiome ^26–30^. However, in general, fibers as therapeutics are not chosen by rational design, selection or optimized to serve a specific function. In this study, we used an *ex vivo* fecal fermentation platform to screen a library of hundreds of novel synthetic and well-characterized glycans ^31^ for propionate production from which we identified KB39 as a high propionate-producing glycan. We also found that *in vivo*, KB39 promotes propionate production and the enrichment of propiogenic bacterial taxa and has beneficial effects on blood lipids and insulin, and preventing liver steatosis and inflammation as well as atherosclerosis in *Ldlr^-/-^* mice fed a high fat, high cholesterol western diet. Our data show that glycans that promote propionate production such as KB39 can provide beneficial effects in lowering multiple CMRFs and preventing cardiometabolic diseases. Our study also underscores the utility of screening large libraries of complex synthetic glycans for identifying novel chemical entities with high therapeutic potential.

## METHODS

### Human fecal sample collection and fecal slurry preparation

Informed consent was obtained from all donors before fecal sample collection. T2DM subjects were males and females between 18 and 70 years old with a BMI between 25 and 45 kg/m^2^ and currently under treatment by a physician for T2DM. Fecal samples were collected into stool specimen containers, immediately frozen and later stored at −80 °C. To prepare fecal slurries, samples were transferred into an anaerobic chamber (Coy) and placed into filtered blender bags (Interscience). Samples were then diluted in 1x PBS and glycerol to a final 20% w/v fecal slurry containing 15% glycerol. Diluted samples were homogenized in a lab blender (Interscience 032230), aliquoted, flash frozen in a dry ice/ethanol bath and stored at −80 °C.

### Microbial cultivation

20% fecal slurry aliquots were transferred into an anaerobic chamber (Coy), thawed and further diluted to a final 1% (w/v) using *Clostridium* minimal medium ^32^ supplemented with 0.1% (w/v) trypticase peptone and 0.75 mM urea. 1% fecal suspensions were dispensed into the wells of 96-well deep well plates containing 5 g/L (w/v) of the appropriate glycan or water used as a negative control. Each treatment was tested in triplicates. Fecal microbial cultures were incubated anaerobically at 37 °C for 45 hours. Plates were then centrifuged at 3,000 x *g* for 10 min at 4 °C. Culture supernatants were collected and stored at −80 °C for acetate, butyrate and propionate quantification. The plates containing fecal bacterial cell pellets were stored at −80 °C for DNA extraction and shallow shotgun sequencing.

### Synthesis of KB39

D-(+)-Galactose (60.0 g, 0.333 mole), D-(+)-mannose (40.0 g, 0.222 mol), citric acid (3.27 g, 0.017 mol, 3 mol%), and deionized water (10 mL) were added to 1 L three-neck round-bottom flask. The flask was equipped with a heating mantle configured with an overhead stirrer, a probe thermocouple, and a reflux condenser in a distillation position to remove excess water throughout the course of the reaction. The temperature controller was set to 130 °C and stirring was initiated as the temperature of the reaction mixture was brought to 130 °C under ambient pressure. The resulting solution was maintained at 130 °C for about 3.5 hours at which time most of the water was removed by distillation and the reaction was determined to be complete by SEC. The heat was turned off and the crude reaction mixture was allowed to cool to <80 °C while maintaining constant stirring. The crude reaction mixture was diluted with deionized water (ca. 60 mL) to obtain a 40 °Bx solution. The resulting solution was passed through a cationic exchange resin (Dowex^®^ Monosphere 88H) column, an anionic exchange resin (Dowex^®^ Monosphere 77WBA) column, and a decolorizing polymer resin (Dowex^®^ OptiPore SD-2) column. The resulting aqueous solution of KB39 was adjusted to ~20 °Bx and then freeze-dried to afford the title compound that could be used without additional purification. The KB39 samples used for *ex vivo* experiments were further purified by chromatography using ISCO RediSep Gold Amine column on a Biotage Isolera equipped with an ELSD detector and water/acetonitrile as mobile phase.

### Glycan linkage analysis

Permethylation was performed as previously described with slight modification ^33^. The glycan sample (500 μg) was dissolved in DMSO for 30 min with gentle stirring. A freshly prepared sodium hydroxide suspension in DMSO was added, followed by a 10 min incubation. Iodomethane (100 μL) was added, followed by a 20 min incubation. A repeated round of sodium hydroxide and iodomethane treatment was performed for complete permethylation. The permethylated sample was extracted, washed with dichloromethane (DCM), and blow dried with nitrogen gas. The sample was hydrolyzed (2M TFA for 2h), reduced with sodium borodeuteride (10 mg mL-1 in 1M ammonia for overnight), and acetylated using acetic anhydride/TFA. The derivatized material was extracted, washed with DCM, and concentrated to 200 μL. Glycosyl linkage analysis was performed on an Agilent 7890A GC equipped with a 5975C MSD detector (EI mode with 70 eV), using a 30-meter RESTEK RTX^®^-2330 capillary column. The GC temperature program: 80 °C for 2 min, a ramp of 30 °C min-1 to 170 °C, a ramp of 4 °C min-1 to 245 °C, and a final holding time of 5 min. The helium flow rate was 1 mL/min, and the sample injection was 1 μL with a split ratio of 10:1.

### DNA sequencing of fecal bacterial cell pellets, mouse fecal samples and microbiome analysis

The DNA from the fecal bacterial pellets and mouse fecal samples were characterized using a shallow shotgun sequencing approach like the one previously described ^34^. Briefly, fecal samples’ DNA was extracted with Qiagen’s DNeasy PowerSoil, quantified using the Quant-iT PicoGreen dsDNA assay (Thermo Fisher) and libraries prepared using the NexteraXT kit and sequenced using HiSeq 1 × 150-cycle v3 kit (Illumina). The operational taxonomic unit (OTU) count tables were generated from the DNA sequences using the SHOGUN pipeline by aligning the filtered reads to a curated database containing all representative genomes in RefSeq for bacteria with additional manually curated strains (Venti) using fully gapped alignment with BURST (https://github.com/knights-lab/BURST) ^34^. The generated OTUs that could be resolved to the species level and the samples that were retained after rarefying to 10,000 reads without replacement were used for downstream analyses. For functional profiling, metagenomic samples were profiled using HUMAnN2 v2.8.2 with default settings ^35^. Gene family files were regrouped using the humann2_regroup_table command with ‘uniref90_ko’ and renormalized to relative abundance using humann2_renorm_table including reads unmapped to strains and unintegrated to pathways. Taxa with relative abundances greater than or equal to 0.1% in either treatment group were retained for downstream analysis. KO (KEGG Orthology) families identified were retained for further analysis if (1) they were among the top 100 terms in at least one sample and (2) they were associated with microbial synthesis of propionate ^36^. To evaluate the treatment effect on the overall microbiome structure, we calculated pair-wise Bray-Curtis dissimilarities based on square-root transformed relative abundances of bacterial species. We then performed principal coordinate analysis on the dissimilarity matrix. To identify taxa and KO families responding to KB39 treatment *ex vivo*, we performed paired Wilcoxon rank sum test between the KB39 treated samples and water treated samples across different communities. For the mouse *in vivo* study, we assessed whether the abundance of a taxon/KO term significantly differed between the KB39 treatment group and the no treatment group at the end of the study using unpaired Wilcoxon rank sum test. The test results were corrected using FDR.

### Short-chain fatty acids quantification

Acetate, butyrate and propionate in microbial culture supernatants were quantified by gas chromatography with flame-ionization detection (GC-FID) (7890A, Agilent). For each sample and calibration standards, 50 μL was mixed with 20 μL of a 400mM 2-ethylbutyric acid (used as an internal standard) solution prepared in HPLC water. Immediately before injection, each sample was acidified with 20 μL of 6% formic acid and 1 μL injected. The GC-FID conditions used were as follows: 15 m x 0.53 mm x 0.50 μm DB-FFAP column (Agilent) - carrier gas: helium at 28.819 mL/min - inlet conditions: 250 °C, 5 mL/min purge flow, 4:1 split ratio – oven temperature gradient: 70 °C, increase at a rate of 70 °C for 1 min, increase at a rate of 100 °C for 1 min, 1.8 min hold. Acetate, butyrate, propionate and succinate in mouse cecal content samples were quantified by liquid chromatography with tandem mass spectrometry (LC-MS/MS). Briefly, 4% (w/v) mouse cecal content homogenates were prepared in 1:1 (v/v) acetonitrile (ACN):water by vortex mixing for 5 min and sonicating for 5 min to extract analytes. Homogenates were then further diluted to 1% (w/v) in 1:1 (v/v) ACN:water, centrifuged and supernatants collected. Supernatants were derivatized in 40 mM 3-nitrophenylhydrazine (3NPH), 37.5 mM N-(3-dimethylaminopropyl)-N’-ethylcarbodiimide hydrochloride, and 1.5% (v/v) pyridine in 1:1 v/v ACN:water at 40 °C for 90 min. The derivatized samples were then further diluted with 10% ACN in water and spiked with labeled internal standards (reacted similarly) before LC-MS/MS analysis. All standards were purchased from Millipore-Sigma. For the HPLC, a Kinetex 2.6 μm C18 50 x 2.1 mm column was used and 0.01% formic acid in water and 0.01% formic acid in methanol were used as mobile phase A and mobile phase B respectively. The MS/MS was run in positive mode and the transitions used were as follows: acetic acid-3NPH: 196.33>136.99 – butyric acid-3NPH: 224.35>137.38 – propionic acid-3NPH: 210.34>137.29 – succinic acid-3NPH: 389.35>137.11. The total amount of each SCFA was calculated based on the weight of the cecum content for each animal.

### Animals and treatment

Nine- to 11-week-old male C57BL/6J *Ldlr^-/-^* mice (Jackson Laboratory, *B6.129S7-Ldlr^tm1Her^/J*, Stock No: 002207) were split into one of three treatment groups (12 animals per group) where they received treatment for 16 weeks: 1) no treatment, western diet only; 2) western diet plus KB39; 3) western diet plus fenofibrate. The western diet consisted of 40 kcal% fat and 1.5% cholesterol (Research Diets D12079B). Mice in the KB39 group were fed the western diet supplemented with KB39 at 7.5% w/w. The KB39 treatment diets were modified to provide the same caloric value to the mice as those mice in the no treatment group. Mice in the fenofibrate group were fed the western diet supplemented with fenofibrate (100 mg/kg/day). Mice in each treatment group received one week of normal chow before the study began. The body weight for each animal was measured before the treatment period and weekly over the treatment period. Four-day food intake for each cage was measured weekly over the treatment period and averaged as food intake per mouse per day.

### Mouse stool, blood and tissue

Fresh fecal samples were collected within a week before treatment initiation and during week 15 post treatment initiation. For each fecal collection period, one fresh fecal pellet was collected on three separate days for each animal, frozen on dry ice and stored at −80 °C. At week 12, an oral glucose tolerance test was performed. Blood was collected from a tail cut after a 6h fast and baseline glucose measured using a glucometer (Lifescan, Johnson & Johnson). The mice then received 2 g/kg of a 100 mg/mL glucose solution in sterile water delivered by oral gavage and blood was collected from the tail wound at 15, 30, 60 and 120 min for glucose measurement. On the day of termination, and an overnight fast, retroorbital blood was collected under anesthesia for the quantification of serum total cholesterol (colorimetric assay, WAKO Diagnostics), serum very low-density lipoprotein (VLDL), serum low-density lipoprotein (LDL) and serum high-density lipoprotein (HDL) (Lipoprint, Qantimetrix), serum non-esterified fatty acids (NEFA) (colorimetric assay, WAKO Diagnostics), serum insulin (MA2400 Mouse/Rat insulin kit K152BZC, Meso Scale Discovery) and whole blood triglycerides (Cardiochek, PTS Diagnostics). The animals were then euthanized, their cecum content collected, and their whole liver dissected and weighed. Liver sections (~100 mg each) were collected and frozen for the quantification of hepatic triglycerides and hepatic total cholesterol. One liver lobe was also preserved in 10% neutral buffered formalin (NBF) for embedding, sectioning and staining with H&E before being scored for severity of steatosis, lobular inflammation and hepatocellular ballooning ^37^. The NAFLD activity score (NAS) was calculated as the sum of liver steatosis, lobular inflammation, and hepatocellular ballooning. The aortic arch was perfused with PBS, dissected, fixed in 10% NBF and later surface stained with oil red O for the quantification of atherosclerotic plaque surface area using the ImagePro Plus software. The heart was dissected, fixed in 10% NBF for embedding and sectioning. Sections of the aortic sinus were stained with elastin trichrome and the atherosclerotic plaque area quantified by morphometric analysis using the ImagePro Plus software. In addition, the foamy macrophages, extracellular lipids and plaque fibrosis in the aortic sinus sections were scored blindly (0: no change; 1: minimal change; 2: mild change, distinct but not prominent; 3: moderate change, prominent tissue feature not effacing pre-existing structures; 4: marked to severe change, overwhelming tissue feature that may efface pre-existing tissue features) and the overall plaque severity score calculated as the sum of the three.

### Hepatic gene expression analysis

Gene expression in liver tissue was analyzed by RNAseq. Total RNA was extracted and RNA samples quantified using Qubit 2.0 Fluorometer (Life Technologies, Carlsbad, CA, USA) and RNA integrity was checked using Agilent TapeStation 4200 (Agilent Technologies, Palo Alto, CA, USA). RNA sequencing libraries were prepared using the NEBNext Ultra RNA Library Prep Kit for Illumina following manufacturer’s instructions (NEB, Ipswich, MA, USA). The sequencing libraries were validated on the Agilent TapeStation (Agilent Technologies, Palo Alto, CA, USA), and quantified by using Qubit 2.0 Fluorometer (Invitrogen, Carlsbad, CA) as well as by quantitative PCR (KAPA Biosystems, Wilmington, MA, USA). The sequencing libraries were pooled and sequenced using a 2×150bp Paired End (PE) configuration using a HiSeq 4000. Raw sequence data files were converted into fastq files and de-multiplexed using Illumina’s bcl2fastq 2.17 software. One mismatch was allowed for index sequence identification. After investigating the quality of the raw data, sequence reads were trimmed to remove possible adapter sequences and nucleotides with poor quality using Trimmomatic v.0.36. The trimmed reads were mapped to the murine reference genome available on ENSEMBL using the STAR aligner v.2.5.2b. Unique gene hit counts were calculated by using feature Counts from the Subread package v.1.5.2. Unique reads that fall within exon regions were counted. After extraction of gene hit counts, the gene hit counts table were used for downstream differential expression analysis. Using DESeq2 ^38^, a comparison of gene expression between the groups of samples was performed. The Wald test was used to generate *P*-values and Log2 fold changes. A principal component analysis (PCA) was performed on log transformed count per million values (logCPM) to compare the treatment impact of KB39 or fenofibrate compared to no treatment on the transcriptomics profile.

## RESULTS

### Screening of a library of synthetic glycans and identification of KB39 as a high propionate-producing glycan

A library of hundreds of synthetic glycans with diverse monosaccharide composition, bond type, branching, and degrees of polymerization was produced as previously described ^31^, of which 593 compounds (Figure S1) were screened for their ability to promote propionate production in a semi-automated *ex vivo* fermentation assay using the fecal microbiota from a healthy human subject. While propionate could be identified in the culture supernatants, there was a wide-range of concentrations, with some glycans supporting 10-fold higher propionate production than others (Figure 1A). This *ex vivo* screen revealed one synthetic glycan, KB39, to be among the glycans enabling the highest propionate production, increasing propionate by 14.5 mM compared to the negative control (Figure 1A, red bar vs blue bar). KB39 (average MW = 2260 g/mol) is composed of D-galactose and D-mannose (60/40 ratio) with complex distribution of glycosidic bonds linking the monosaccharides (Figure S2). The linkage and extensive branching contrast with known naturally occurring dietary fibers ^31^.

**Figure 1.**
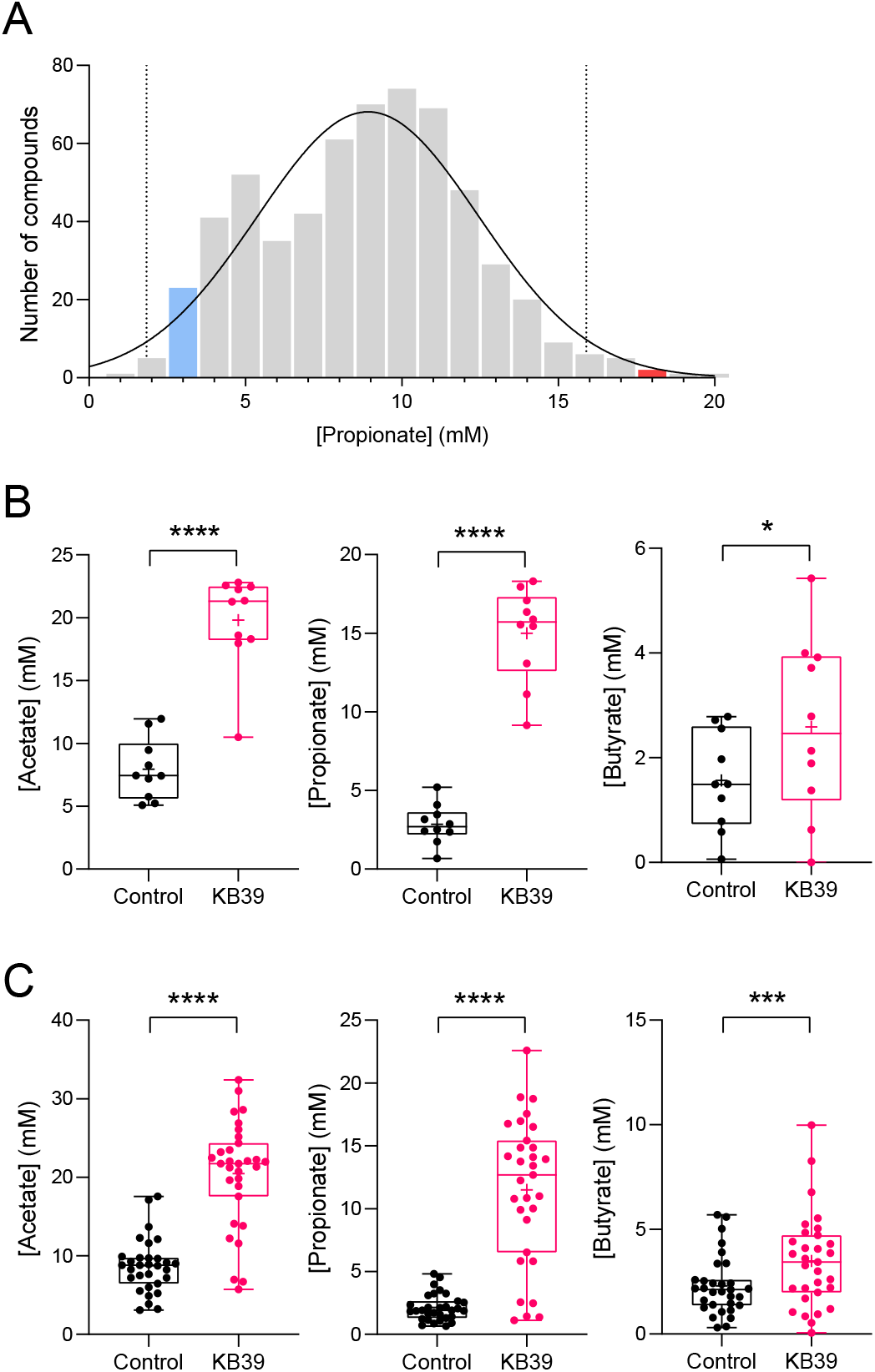
Identification of high propionate-producing synthetic glycan KB39. **(A)** High-throughput *ex vivo* screen of synthetic glycan library with fecal microbiota from a healthy subject. Fecal microbiota cultures were incubated with synthetic glycans under anaerobic conditions and propionate was measured in culture supernatants by gas chromatography with flame ionization detection (GC-FID). Data represent the distribution of propionate production. The blue and red bars indicate where the no added carbon control and KB39 are on the distribution, respectively. The curve represents the normal distribution and the dotted lines indicate the 95% confidence interval. **(B, C)** Production of each SCFA by KB39 with fecal microbiota from 10 healthy subjects (B) and 31 overweight-T2DM patients (C). Fecal microbiota cultures were incubated without (negative control) or with KB39 and SCFA were measured in culture supernatants. * *P*<0.05, *** *P*<0.001**** *P*<0.0001, paired *t*-test.

The composition and metabolic function of gut microbiota can vary across individuals. To evaluate the reproducibility of propionate production by different gut microbiota, we tested KB39 in our *ex vivo* assay system with fecal microbiota samples from ten healthy subjects and 31 overweight subjects with diagnosed T2DM, a population reported to have altered gut microbiomes ^18^. High propionate production was observed for most fecal samples incubated with KB39 and was similar between healthy and T2DM subjects (Figure 1B vs. 1C, center panels). The fermentation of KB39 with these fecal samples also resulted in a similar increase in acetate production, and a more moderate increase in butyrate (Figure 1B, 1C). Importantly, KB39 increased propionate fraction of total SCFAs, unlike acetate and butyrate (Figure S3). This fermentation product selectivity suggests that KB39 was disproportionately metabolized by propionate-producing bacteria.

### KB39 enriches the propionate-producing phylum *Bacteroidetes* in fecal microbiota from overweight-T2DM subjects *ex vivo*

The increase and disproportionate production of propionate by fecal microbiota incubated with KB39 suggested that it preferentially supported the growth of propiogenic commensal bacteria. Although the ability to produce propionate is shared by numerous bacterial species in the gut, it is particularly widespread among the species in the phylum *Bacteroidetes*. To identify the commensal bacteria primarily responsible for KB39 fermentation in samples from overweight-T2DM patients, we subjected the *ex vivo* fecal microbiota cultures to metagenomics sequencing reasoning that the bacteria responsible for KB39 fermentation would increase in relative abundance compared to the rest of the bacteria in the cultures. We observed a significant enrichment of the phylum *Bacteroidetes* (7-fold increase), which was driven primarily by an increase in the relative abundance of several species belonging to the genus *Parabacteroides* (51-fold increase) in the family *Tannerellaceae* (Figure 2). We also observed a lower magnitude enrichment of *Bacteroides fragilis* and *Bacteroides thetaiotaomicron* species. Thus, KB39 fermentation and the production of propionate likely derives from the growth and metabolic activity of species of *Parabacteroides* and *Bacteroides*. In addition, metagenomic sequencing also revealed an increase in the relative abundance of *Eisenbergiella tayi*, a species in the family *Lachnospiraceae* known to produce butyrate as its main fermentation product, in addition to lactate, acetate and succinate ^39^. Furthermore, significant increases in the relative abundance of the family *Erysipelotrichaceae* and the species *Blautia producta* were also detected. The family *Erysipelotrichaceae* is commonly found in the human gut microbiome and contains the genera *Holdemania* and *Turicibacter* known to produce acetate and lactate, respectively ^40, 41^. *Blautia producta* produces lactate and acetate as its main products of fermentation ^42^. By contrast, KB39 decreased the relative abundance of *Citrobacter, Escherichia coli, Enterococcus faecalis* and *Klebsiella pneumoniae*, bacterial species generally considered as opportunistic pathogens and pro-inflammatory.

**Figure 2.**
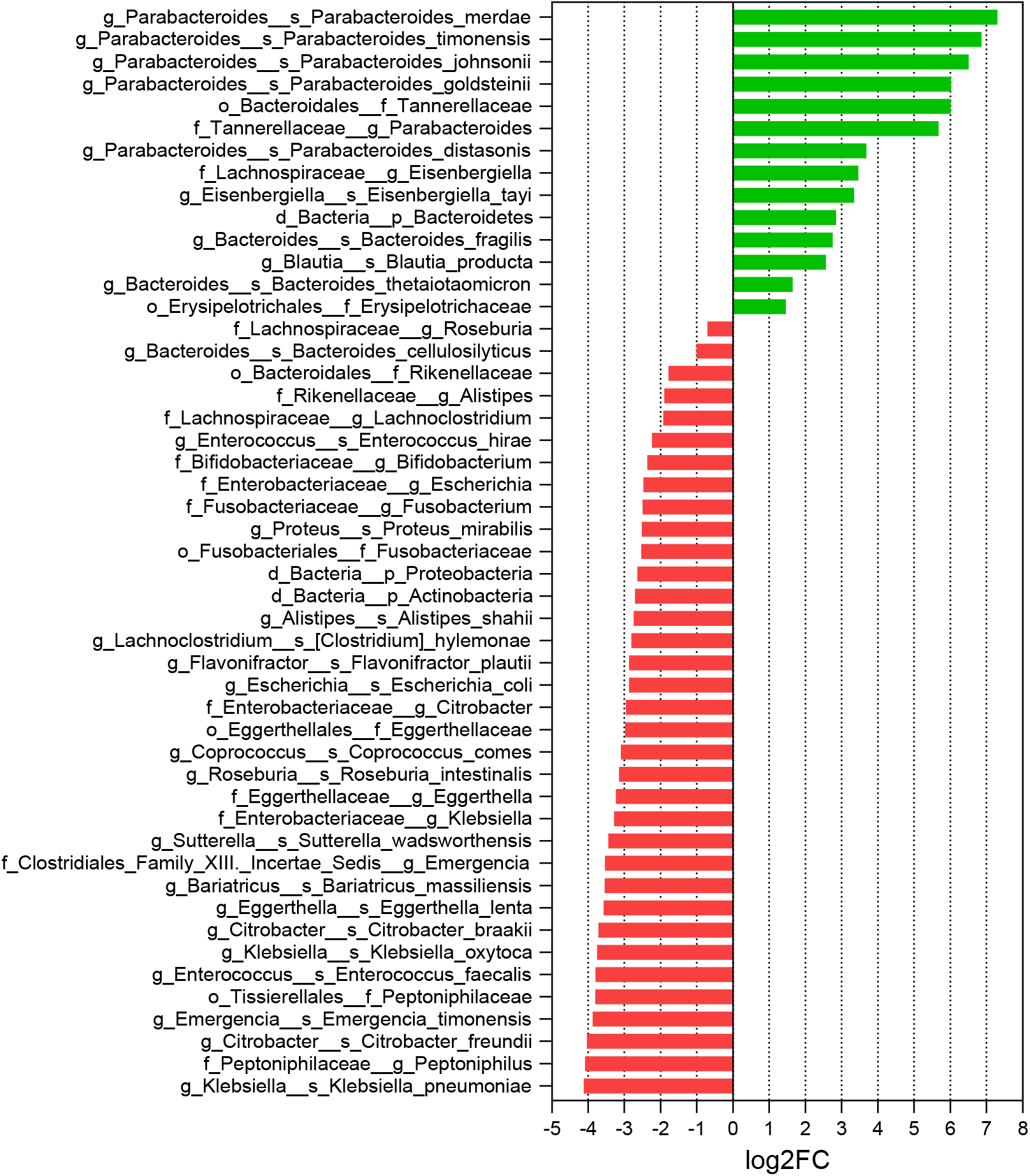
KB39 enriches propionate-producing commensal bacteria *ex vivo* in overweight-T2DM subjects. Fecal microbiota cultures from 31 overweight-T2DM subjects were incubated *ex vivo* without (negative control) or with KB39. Significantly enriched and depleted taxa (fdr < 0.1, Wilcox rank sum test with FDR correction) at phylum, family, genus, and species level in KB39 treated group in comparison to water. Data represent the log2 fold change.

### KB39 increases cecal propionate and propionate-producing *Bacteroidetes in vivo* in western diet-fed *Ldlr^-/-^* mice

The effect of KB39 on propionate production *in vivo* was determined in LDL receptor knockout (*Ldlr^-/-^*) mice fed a western diet (high fat, high cholesterol), a model known to develop multiple cardiometabolic risk factors (CMRFs) such as obesity, hypercholesterolemia, hyperinsulinemia, mild hyperglycemia and insulin resistance resulting in fatty liver and atherosclerosis ^43–45^. Briefly, *Ldlr^-/-^* mice were fed a western diet supplemented with KB39 or fenofibrate for 16 weeks. Fenofibrate, a peroxisome proliferator receptor alpha (PPARα) agonist, was used as a positive control for its documented anti-obesity, lipid and glucose lowering effects in the western diet-fed *Ldlr^-/-^* mouse model ^46, 47^. Similar to what was observed *ex vivo* in fecal communities from humans, treatment with KB39 increased the levels of total cecal acetate and propionate compared to no treatment (Figure 3A). A small increase in cecal butyrate was also observed, although non-significant. In addition, KB39 decreased the proportion of acetate and butyrate and increased the proportion of cecal propionate compared to no treatment, similar to what was observed *ex vivo* (Figure 3B) and suggesting an enrichment in propiogenic bacteria *in vivo*. KB39 also increased the total amount and proportion of succinate (Figure 3A, 3B). The “succinate pathway” of propionate formation is the major propionate biosynthetic pathway in the large intestine for taxa of the *Bacteroidetes* phylum ^36^. This result suggests that KB39 is converted to propionate via the succinate pathway *in vivo* and that succinate might accumulate due to limiting factors such as vitamin B12 ^48^. As expected, fenofibrate did not modify the total amount or proportion of SCFAs or succinate (Figure 3A, 3B).

**Figure 3.**
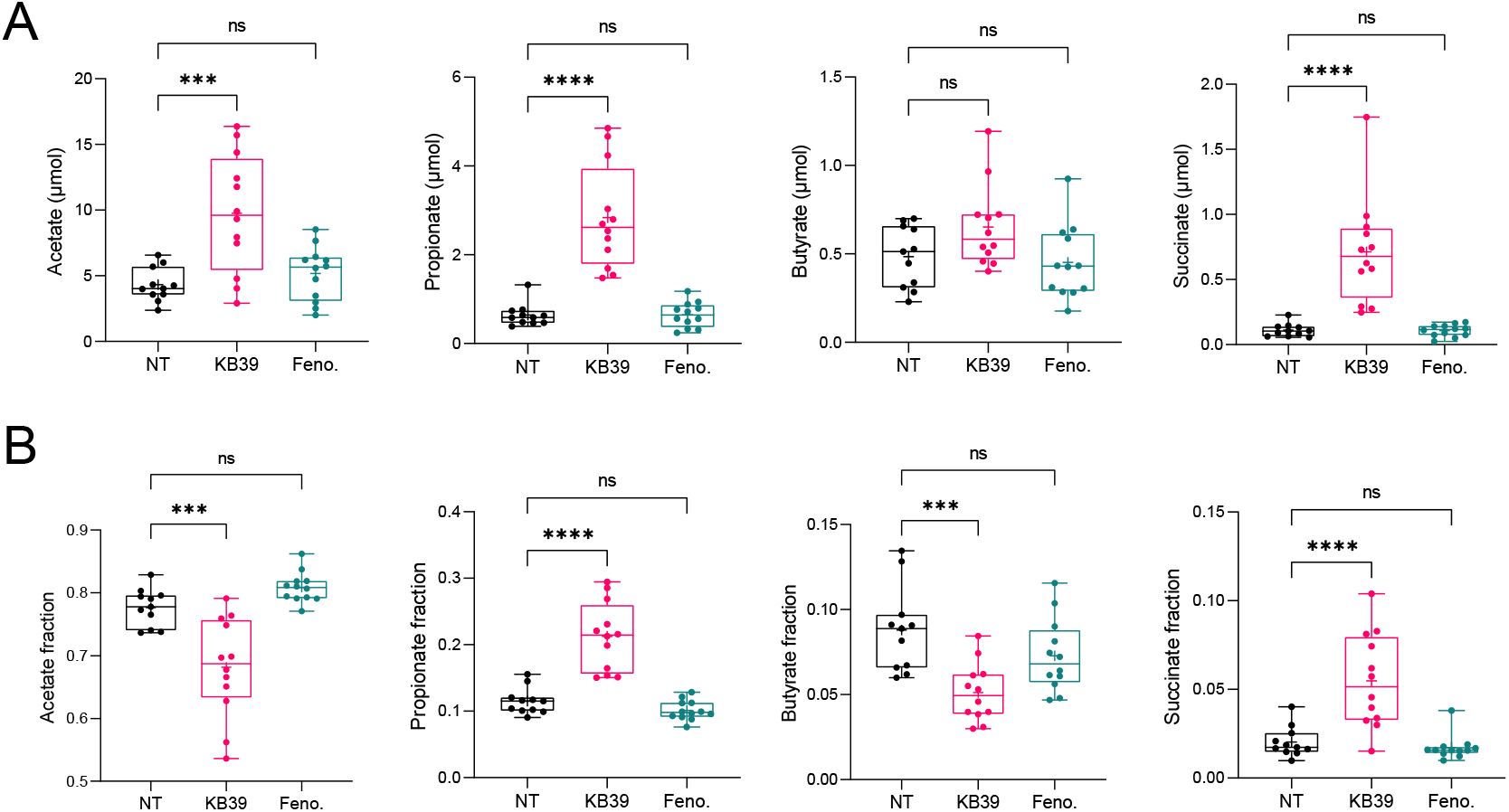
KB39 increases propionate, acetate and succinate production and increases the proportion of cecal propionate and succinate in the cecum of western diet-fed *Ldlr^-/-^* mice. Male *Ldlr^-/-^* mice were fed a western diet, western diet supplemented with KB39 (7.5% w/w) or fenofibrate (100 mg/kg/day) for 16 weeks. **(A)** Total cecal acetate, butyrate, propionate and succinate. **(B)** Acetate, butyrate, propionate and succinate relative fractions. NT: no treatment; Feno: fenofibrate. *** *P*<0.001, **** *P*<0.0001, 1-way ANOVA with Dunnett pairwise comparison to NT group.

The effect of KB39 on the composition of the gut microbiome was determined by shotgun sequencing of stool samples collected before and at the end of the treatment period. The principal coordinate analysis (PCoA) plot of the microbiome composition data shown in Figure S4 shows that the western diet had a profound impact on the fecal microbiome composition of *Ldlr^-/-^* mice. Western diet feeding led to a decrease in major SCFA producing taxa, including *Bacteroidetes* and *Ruminococcaceae*, and the reduction in *Akkermansia muciniphila*, a species associated with reduced obesity and type 2 diabetes ^49^, in non-treated animals (Figure S5). Unlike fenofibrate, treatment with KB39 led to a widely distinct microbiome structure (Figure S4). Compared to no treatment, KB39 enriched propiogenic families and genera belonging to the *Bacteroidetes* phylum, including *Rickenellaceae, Tannerellaceae, Alistipes* and *Parabacteroides* (Figure 4). As noted above, this enrichment in propionate producing *Bacteroidetes*, which are known to utilize the “succinate pathway” for propionate formation, is consistent with the *ex vivo* data and the observed increase in cecal succinate in KB39-treated animals compared to no treatment. This was corroborated by the increase, both *ex vivo* and *in vivo*, in the abundance of sequencing reads mapping to genes encoding the methylmalonyl-CoA epimerase enzyme, essential for propionate production via the “succinate pathway” and used by *Bacteroidetes* bacterial species (Figure S6) and no response of genes associated with the two other (acrylate and propanediol) propionate synthesis pathways ^36^. KB39 also promoted expansion of the acetate producer *Blautia producta* and depletion of *Enterococcus faecalis*, consistent with what was observed *ex vivo* with human fecal gut microbiota. Notably, KB39 also enriched *Akkermansia muciniphila* mentioned previously, and *Parasutterella*, a genus belonging to the *Proteobacteria* phylum and recently shown to be associated with reduced LDL-C in humans ^50^.

**Figure 4.**
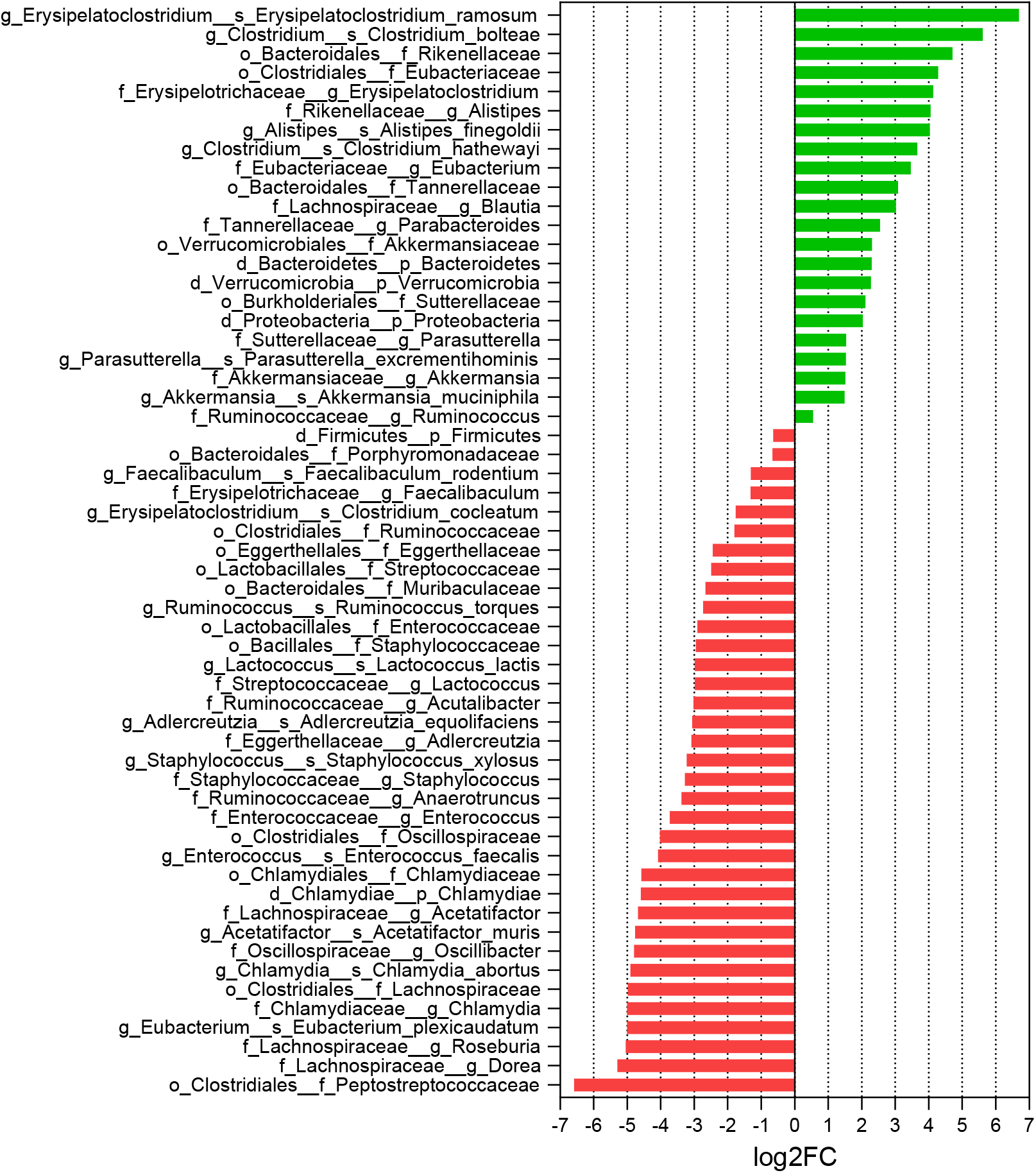
KB39 shifts the gut microbiome in western diet-fed *Ldlr^-/-^* mice and promotes propionate-producing bacterial taxa. Male *Ldlr^-/-^* mice were fed a western diet, western diet supplemented with KB39 (7.5% w/w) or fenofibrate (100 mg/kg/day) for 16 weeks. Fecal samples were collected before treatment during the last week of the treatment period. Significantly enriched and depleted taxa (fdr < 0.1, Wilcox rank sum test with FDR correction) at phylum, family, genus, and species levels in KB39 treated group in comparison to no treatment group at the end of the study.

### KB39 attenuates hypercholesterolemia, hyperinsulinemia in WD-fed *Ldlr^-/-^* mice

Following on the confirmation that KB39 increases propionate and propionate-producing gut bacteria *in vivo*, we evaluated the effect of KB39 on the metabolic parameters of western diet fed *Ldlr^-/-^* mice. Unlike fenofibrate that dramatically prevented body weight gain, KB39 only had a modest effect on relative body weight gain during the treatment period (Figure 5A), and no effect on food intake (Figure 5B). However, KB39 treatment greatly lowered serum total cholesterol (43%), LDL-C (39%), VLDL-C (58%), and HDL-C (47%) and provided a lowering trend for non-esterified fatty acids (NEFA) (16%) compared with no treatment (Table 1). By contrast, treatment with fenofibrate provided non-significant lipid lowering trends except for a decrease in serum LDL-C (23%) and NEFA (41%) (Table 1). In addition to the large decrease in serum cholesterol, KB39 also significantly reduced hepatic total cholesterol (39%) and hepatic triglycerides (37%) compared with no treatment. Fenofibrate appeared to lead to a similar decrease in hepatic cholesterol (33%) as KB39 but provided a larger decrease in hepatic triglycerides (73%). The effect of KB39 on liver fat was accompanied by lower liver weights (Table 1). By contrast, consistent with previous reports of hepatomegaly and peroxisome proliferation in response to fibrates in mice ^51^, liver weights in mice treated with fenofibrate significantly increased, despite reductions in hepatic cholesterol and triglyceride levels. Both KB39 and fenofibrate decreased fasting serum insulin levels by ~66%, consistent with reduced insulin resistance, but only fenofibrate significantly reduced fasting blood glucose levels or improved glucose tolerance during an OGTT (Table 1, Figure 5C). Taken together, these data show that KB39 reduces multiple CMRFs with, in the cases of cholesterol and insulin, effect sizes comparable to fenofibrate, a compound used to treat high cholesterol and triglycerides.

**Figure 5.**
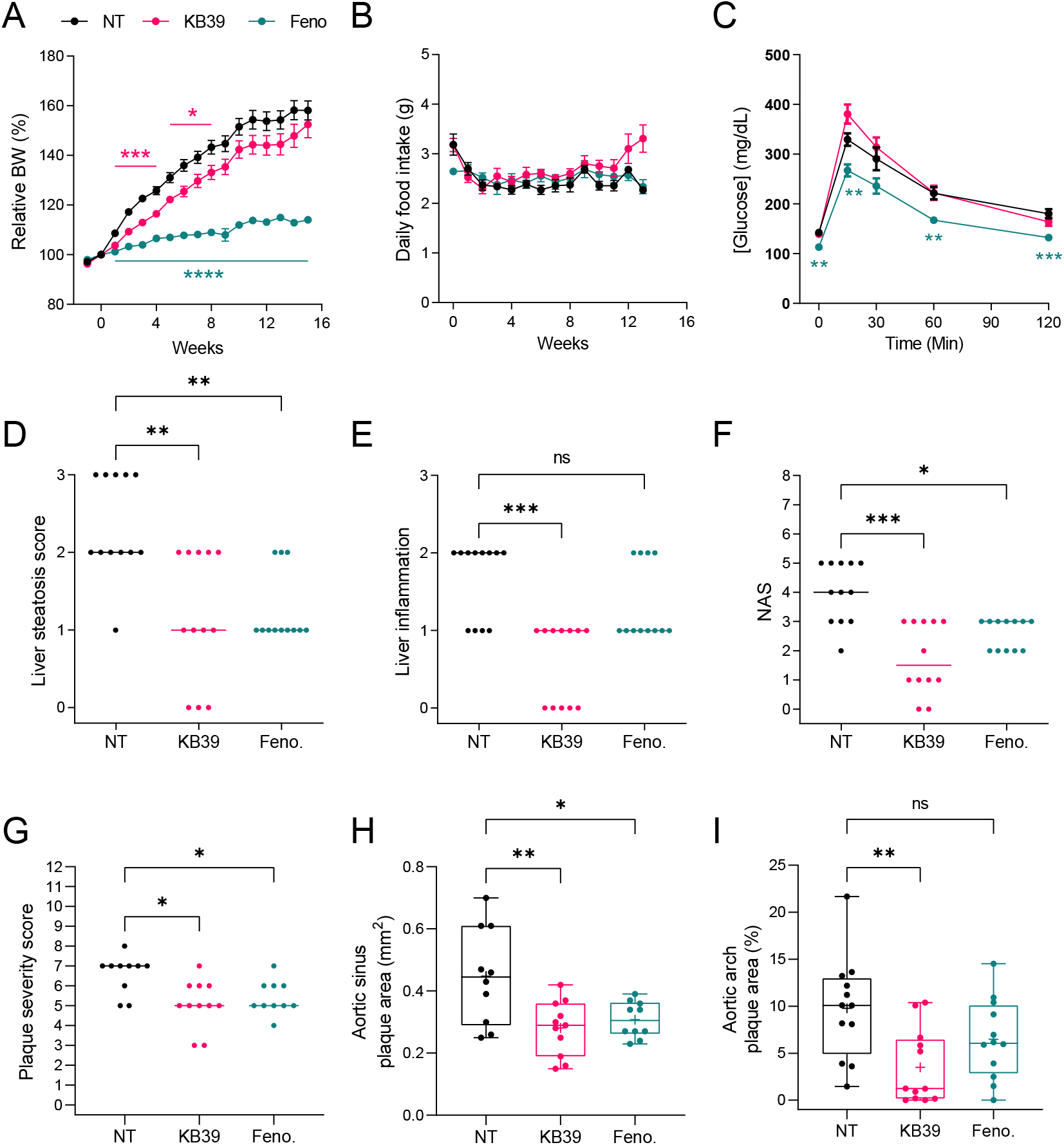
KB39 attenuates liver steatosis, liver inflammation and atherosclerosis in western diet-fed *Ldlr^-/-^* mice. Male *Ldlr^-/-^* mice were fed a western diet, western diet supplemented with KB39 (7.5% w/w) or fenofibrate (100mg/kg/day) for 16 weeks. An OGTT was performed at week 15. **(A-C)** Relative body weight (BW) compared to BW at treatment initiation (A), daily food intake (B) and OGTT blood glucose measurements (C); mean ± SEM, 2-way ANOVA, Dunnett. **(D-F)** Liver steatosis score (D), inflammation (E) score and NAFLD activity score (NAS) (F) at termination; median, Kruskal-Wallis, Dunn. **(G)** Aortic sinus plaque severity score; median; Kruskal-Wallis, Dunn. **(H, I)** Plaque area in the aortic sinus quantified by morphometric analysis of elastin trichrome-stained sections (H) and in the aortic arch quantified by oil red O staining “en face” analysis (I), 1-way ANOVA, Dunnett. * *P*<0.05, ** *P*<0.01, *** *P*<0.001, **** *P*<0.0001. NT: no treatment; Feno: fenofibrate.

**Table 1.** Effects of KB39 and fenofibrate in western diet-fed *Ldlr^-/-^* mice. Male *Ldlr^-/-^* mice were fed a western diet, western diet supplemented with KB39 (7.5% w/w) or western diet supplemented with fenofibrate (100 mg/kg/day) for 16 weeks. An OGTT was performed at week 15. Measurements were taken in fasted animals; see material and methods for details. Data represent the mean ± SEM; statistics: 1-way ANOVA with Dunnett pairwise comparison to no treatment group.

|  | Mean ± SEM |  |  | P-value |  |
| --- | --- | --- | --- | --- | --- |
|  | No Treatment | KB39 | Fenofibrate | KB39 | Fenofibrate |
| Serum total cholesterol (mg/dL) | 1924.3±140.4 | **1094.7±132.4** | 1564.3±59.6 | **<0.0001** | 0.0660 |
| Serum VLDL (mg/dL) | 449.9±63.0 | **190.9±39.6** | 344.4±46.6 | **0.0019** | 0.2457 |
| Serum LDL (mg/dL) | 949.5±76.8 | **581.1±66.1** | **728.4±36.2** | **0.0004** | **0.0298** |
| Serum HDL (mg/dL) | 601.5±73.4 | **321.3±47.4** | 484.9±36.9 | **0.0016** | 0.2230 |
| Blood triglycerides (mg/dL) | 354.9±28.3 | 293.8±24.1 | 317.3±29.1 | 0.2105 | 0.5243 |
| Serum NEFA (mM) | 1.11±0.07 | 0.93±0.05 | **0.65±0.05** | 0.0683 | **<0.0001** |
| Hepatic total cholesterol (mg/g) | 12.61±0.84 | **7.66±0.48** | **8.50±0.31** | **<0.0001** | **<0.0001** |
| Hepatic triglycerides (mg/g) | 156.8±13.5 | **98.6±10.2** | **42.4±5.4** | **0.0006** | **<0.0001** |
| Blood glucose (mg/dL) | 142.1±5.7 | 139.4±6.8 | **113.1±5.1** | 0.9284 | **0.0028** |
| Blood glucose AUC (g/dL.min) (OGTT) | 10.9±3.5 | 12.0±4.1 | 8.1±2.0 | 0.5909 | 0.0882 |
| Serum insulin (ng/mL) | 1.01±0.18 | **0.35±0.08** | **0.33±0.10** | **0.0022** | **0.0015** |
| Final body weight (g) | 39.3±1.4 | 38.5±1.4 | **29.2±0.6** | 0.8285 | **<0.0001** |
| Body weight change (%) | 58.2±3.8 | 52.5±5.3 | **14.0±1.2** | 0.4800 | **<0.0001** |
| Liver weight (mg) | 1690±112.7 | **1338±81.1** | **2442±81.8** | **0.0218** | **<0.0001** |
| Relative liver weight (% body weight) | 4.27±0.20 | **3.46±0.10** | **8.37±0.27** | **0.0128** | **<0.0001** |

### KB39 attenuates liver steatosis, liver inflammation and atherosclerosis in western diet-fed *Ldlr^-/-^* mice

The beneficial metabolic effects of KB39 in western diet-fed *Ldlr^-/-^* mice reported in Table 1 were associated with improvement in liver histopathology, with decreases in liver steatosis (−1 point), inflammation (−1 point), and overall NAFLD activity score (NAS) (−2.5 points) (Figure 5D, 5E, 5F). Treatment with fenofibrate demonstrated significant decreases in liver steatosis and NAS but had no significant effect on liver inflammation. In addition, consistent with its LDL-C lowering effect, KB39 also had very significant effects on atherosclerosis in this mouse model, with 64% reduction in atherosclerotic plaque formation in the aortic arch (Figure 5I) and a 38% reduction of plaque in the aortic sinus (Figure 5H). KB39 also led to a significant reduction in the overall plaque severity score in the aortic sinus (−2 point) (Figure 5G). Thus, KB39 treatment reduced biochemical markers of cardiometabolic disease risk, reduced fatty liver disease, and produced a major reduction in atherosclerosis development in *Ldlr^-/-^* mice on western diet.

### KB39 upregulates genes involved in fatty acid oxidation and repressed markers of inflammation and insulin resistance in the liver of western diet-fed *Ldlr^-/-^* mice

Given the cholesterol and insulin lowering effects of KB39 described herein and the central role played by the liver in lipid and glucose metabolism, we compared the transcriptional profile of the non-treated, KB39 and fenofibrate-treated western-diet fed *Ldlr^-/-^* mice using RNA-seq. The PCA of liver RNA-seq data presented in Figure S7 shows a distinct liver transcriptional profile between the different groups, confirming the differences observed in liver histology and biochemistry. As shown in Table 2, KB39 significantly increased the expression of PPARα, a central regulator of fatty acid oxidation ^52^, compared to no treatment. Like fenofibrate, a known PPARα agonist, KB39 also increased the expression of genes involved in fatty acid oxidation, including mitochondrial and microsomal oxidation. On the other hand, KB39 significantly repressed the expression of PPARγ and SREBP1 (encoded by *Pparg* and *Srebf1* genes), two major transcriptional regulators involved in hepatic lipogenesis ^52, 53^. Accordingly, *Scd1*, encoding for the stearoyl-coenzyme A desaturase 1, a critical enzyme for the synthesis of monounsaturated fatty acids and triglycerides and positively regulated by PPARγ and SREBP1 ^54–56^, was repressed by KB39 treatment. By contrast, fenofibrate induced the expression of *Srebf1, Scd1* and other major genes involved in fatty acid synthesis and elongation controlled by SREBP1 (*e.g., Acaca, Acacb, Fasn, Elovl6*). These results are consistent with previous studies that showed that PPARα agonists activate both fatty acid oxidation and fatty acid synthesis simultaneously in the liver ^57^. In addition to fatty acid synthesis, KB39 and fenofibrate also differed in their effect on bile acid synthesis and transport genes. KB39 increased the expression of bile acid synthesis genes and the bile acid transporter genes *Abcd11* (*Ntcp*) and *Slc10a1* (*Bsep*). Importantly, KB39 and fenofibrate increased the expression of the master regulator of cholesterol synthesis SREBP2 (encoded by *Srebf2*) and numerous SREBP2-regulated cholesterol synthetic genes compared to no treatment. These results are consistent with the observed decrease in hepatic cholesterol, what is observed with other cholesterol lowering agents such as statins, and the negative transcriptional regulation cholesterol exerts on its own synthesis in the liver ^58, 59^. Notably, and similar to what is observed with statins ^60^, KB39 also increased the expression of the LDL receptor (LDLR). Importantly, in addition to its effect on genes involved in lipid metabolism, KB39 decreased the expression of markers of inflammation, oxidative stress, apoptosis and liver damage. This decrease was associated with a reduction in markers of fibrosis, cell adhesion and macrophage activation, and were consistent with the reduction in hepatic inflammation observed histologically. A downregulation of the markers of insulin resistance *Socs3* and *Trib3* ^61, 62^ was also observed for KB39 treated animals compared to no treatment, supporting an increase in hepatic insulin sensitivity corroborated with the observed decrease in fasting blood insulin. Interestingly, fenofibrate increased the expression of *Trib3*, a known PPARα target with potential negative impacts on hepatic insulin signaling ^63^. KB39 also upregulated *Ceacam1*, an important gene mediating hepatic insulin clearance and the maintenance of insulin sensitivity, known to be downregulated in rodent models of obesity, insulin resistance and NAFLD as well as obese subjects with insulin resistance ^64^. Finally, KB39 increased the expression of multiple markers of autophagy, an important protection mechanism against liver damage and lipotoxicity ^65^.

**Table 2.** Genes differentially expressed in the liver of western diet-fed *Ldlr^-/-^* mice treated with KB39 or fenofibrate. Male *Ldlr^-/-^* mice were fed a western diet, western diet supplemented with KB39 (7.5% w/w) or fenofibrate (100 mg/kg/day) for 16 weeks. Livers were collected at termination and hepatic gene expression was analyzed by RNAseq. Deseq2 ^38^ was used to determine the log2FC and adjusted *P*-value (padj) in gene expression compared to the no treatment group.

| Gene | KB39 |  | Fenofibrate |  | description |
| --- | --- | --- | --- | --- | --- |
|  | log2FC | padj | log2FC | padj |  |
| Fatty acid oxidation |  |  |  |  |  |
| Cpt1a | 0.43 | 1.45e-03 | 0.05 | 6.84e-01 | carnitine palmitoyltransferase 1a |
| Cpt2 | 0.37 | 6.55e-05 | 0.32 | 1.89e-04 | carnitine palmitoyltransferase 2 |
| Hadha | 0.54 | 3.68e-10 | 0.91 | 3.01e-39 | hydroxyacyl-CoA dehydrogenase trifunctional subunit alpha |
| Acads | 0.21 | 9.44E-03 | 0.53 | 2.43E-14 | acyl-Coenzyme A dehydrogenase, short chain |
| Acadm | 0.22 | 1.02E-03 | 0.60 | 7.13E-17 | acyl-Coenzyme A dehydrogenase, medium chain |
| Acadl | 0.22 | 1.04E-03 | 0.87 | 2.17E-31 | acyl-Coenzyme A dehydrogenase, long-chain |
| Hadha | 0.54 | 3.68e-10 | 0.91 | 3.01e-39 | hydroxyacyl-CoA dehydrogenase trifunctional subunit alpha |
| Hadhb | 0.40 | 3.02E-07 | 0.92 | 4.39E-44 | hydroxyacyl-CoA dehydrogenase trifunctional multienzyme complex subunit beta |
| Acaa2 | 0.39 | 1.81e-06 | 0.57 | 6.98e-13 | acetyl-Coenzyme A acyltransferase 2 |
| Abcd3 | 0.38 | 6.03e-06 | 0.71 | 1.27e-10 | ATP-binding cassette, sub-family D (ALD), member 3 |
| Acox1 | 0.32 | 1.91e-03 | 1.54 | 1.24e-49 | acyl-Coenzyme A oxidase 1, palmitoyl |
| Acaa1a | 0.32 | 7.77E-05 | 0.53 | 9.39E-10 | acetyl-Coenzyme A acyltransferase 1A |
| Acaa1b | 0.43 | 2.97e-05 | 1.15 | 4.43e-31 | acetyl-Coenzyme A acyltransferase 1B |
| Ehhadh | 0.58 | 6.02e-02 | 2.98 | 7.56e-105 | enoyl-CoA, hydratase/3-hydroxyacyl Coenzyme A dehydrogenase |
| Cyp4a14 | 1.38 | 2.08e-04 | 2.72 | 1.77e-27 | cytochrome P450, family 4, subfamily a, polypeptide 14 |
| Regulators |  |  |  |  |  |
| Ppara | 0.33 | 5.83e-03 | -0.13 | 3.15e-01 | peroxisome proliferator activated receptor alpha |
| Pparg | -0.60 | 7.49e-04 | -0.63 | 5.73e-06 | peroxisome proliferator activated receptor gamma |
| Srebf1 | -0.43 | 7.04e-03 | 0.98 | 1.02e-25 | sterol regulatory element binding transcription factor 1 |
| Srebf2 | 0.46 | 3.30e-03 | 0.69 | 1.63e-17 | sterol regulatory element binding factor 2 |
| Fatty acid synthesis |  |  |  |  |  |
| Acaca | -0.05 | 7.62e-01 | 1.16 | 7.94e-20 | acetyl-Coenzyme A carboxylase alpha |
| Acacb | 0.37 | 9.00e-02 | 0.96 | 5.46e-09 | acetyl-Coenzyme A carboxylase beta |
| Fasn | 0.11 | 6.24e-01 | 1.98 | 7.25e-27 | fatty acid synthase |
| Elovl6 | 0.11 | 4.42e-01 | 0.47 | 1.27e-04 | ELOVL family member 6, elongation of long chain fatty acids |
| Scd1 | -0.79 | 2.03e-02 | 0.44 | 3.94e-04 | stearoyl-Coenzyme A desaturase 1 |
| Bile acids |  |  |  |  |  |
| Abcb11 (Ntcp) | 0.29 | 1.35e-03 | -0.02 | 8.82e-01 | ATP-binding cassette, sub-family B (MDR/TAP), member 11 |
| Cyp7a1 | 0.69 | 1.61e-03 | -0.13 | 7.00e-01 | cytochrome P450, family 7, subfamily a, polypeptide 1 |
| Cyp7b1 | 1.26 | 4.82e-03 | -0.40 | 2.91e-01 | cytochrome P450, family 7, subfamily b, polypeptide 1 |
| Cyp8b1 | 0.85 | 4.66e-03 | 0.76 | 1.08e-03 | cytochrome P450, family 8, subfamily b, polypeptide 1 |
| Slc10a1 (Bsep) | 0.38 | 8.06e-04 | 0.33 | 4.23e-03 | solute carrier family 10 (sodium/bile acid cotransporter family), member 1 |
| Cholesterol synthesis |  |  |  |  |  |
| Ldlr | 0.25 | 1.35e-02 | 0.13 | 9.70e-02 | low density lipoprotein receptor |
| Hmgcs1 | 0.57 | 1.59e-02 | 1.80 | 1.29e-22 | 3-hydroxy-3-methylglutaryl-Coenzyme A synthase 1 |
| Hmgcr | 0.36 | 8.02e-02 | 0.55 | 2.13e-04 | 3-hydroxy-3-methylglutaryl-Coenzyme A reductase |
| Mvk | 0.46 | 4.23e-02 | 0.85 | 3.72e-08 | mevalonate kinase |
| Mvd | 0.82 | 3.39e-02 | 1.88 | 4.83e-17 | mevalonate (diphospho) decarboxylase |
| Idi1 | 2.22 | 1.59e-04 | 3.04 | 7.36e-34 | isopentenyl-diphosphate delta isomerase |
| Fdps | 0.98 | 2.16e-02 | 2.80 | 8.26e-68 | farnesyl diphosphate synthetase |
| Sqle | 0.64 | 1.93e-01 | 1.92 | 2.89e-10 | squalene epoxidase |
| Cyp51 | 1.53 | 2.14e-04 | 1.84 | 6.71e-26 | cytochrome P450, family 51 |
| Inflammation |  |  |  |  |  |
| Ccl2 (Mcp1) | -2.33 | 1.20e-11 | -1.71 | 1.87e-14 | chemokine (C-C motif) ligand 2 |
| Ccl5 | -1.84 | 2.82e-15 | -1.11 | 3.20e-06 | chemokine (C-C motif) ligand 5 |
| Il1a | -1.03 | 1.67e-08 | -0.50 | 1.00e-03 | interleukin 1 alpha |
| Il1b | -0.64 | 1.21e-02 | -0.38 | 5.06e-02 | interleukin 1 beta |
| Lcn2 | -1.45 | 7.94e-04 | -2.21 | 1.28e-08 | lipocalin 2 |
| Tnf | -2.41 | 1.24e-07 | -1.52 | 3.94e-05 | tumor necrosis factor |
| Oxidative stress |  |  |  |  |  |
| Cybb | -0.99 | 4.04e-10 | -0.64 | 2.54e-06 | cytochrome b-245, beta polypeptide |
| Hmox1 | -1.22 | 1.15e-19 | -0.33 | 1.68e-02 | heme oxygenase 1 |
| Ncf2 | -1.10 | 1.07e-09 | -0.86 | 1.76e-09 | neutrophil cytosolic factor 2 |
| Apoptosis |  |  |  |  |  |
| Bax | -0.33 | 1.09e-02 | -0.20 | 6.06e-02 | BCL2-associated X protein |
| Bcl2 | -0.88 | 2.60e-05 | -0.74 | 6.81e-05 | B cell leukemia/lymphoma 2 |
| Bcl2a1a | -1.25 | 3.03e-06 | -1.10 | 1.35e-06 | B cell leukemia/lymphoma 2 related protein A1a |
| Traf1 | -1.43 | 6.37e-06 | -0.98 | 1.60e-04 | TNF receptor-associated factor 1 |
| Fibrosis |  |  |  |  |  |
| Acta2 (α-Sma) | -1.10 | 3.16e-03 | -0.10 | 8.10e-01 | actin alpha 2, smooth muscle |
| Col1a1 | -2.75 | 1.24e-10 | -1.62 | 6.00e-05 | collagen, type I, alpha 1 |
| Mmp13 | -3.67 | 3.33e-12 | -2.56 | 4.29e-08 | matrix metalloproteinase 13 |
| Tgfb1 | -0.52 | 4.58e-04 | -0.38 | 6.88e-03 | transforming growth factor, beta 1 |
| Timp1 | -3.87 | 1.33e-11 | -2.87 | 6.58e-11 | tissue inhibitor of metalloproteinase 1 |
| Cell adhesion |  |  |  |  |  |
| Icam1 | -0.77 | 1.25e-06 | -0.64 | 3.66e-07 | intercellular adhesion molecule 1 |
| Vcam1 | -1.45 | 1.41e-14 | -0.77 | 5.79e-07 | vascular cell adhesion molecule 1 |
| Kupffer Cell/macrophage |  |  |  |  |  |
| Adgre1 (F4/80) | -0.34 | 7.39e-02 | -0.45 | 8.14e-03 | adhesion G protein-coupled receptor E1 |
| Cd14 | -1.18 | 3.70e-06 | -0.48 | 3.79e-02 | CD14 antigen |
| Cd68 | -1.38 | 1.61e-13 | -0.91 | 5.64e-08 | CD68 antigen |
| Itgam (Cd11b) | -0.93 | 2.73e-03 | -1.09 | 5.38e-05 | integrin alpha M |
| Itgax (Cd11c) | -2.61 | 5.72e-13 | -1.37 | 2.02e-09 | integrin alpha X |
| Insulin signaling |  |  |  |  |  |
| Ceacam1 | 0.48 | 1.78e-05 | 0.08 | 4.46e-01 | carcinoembryonic antigen-related cell adhesion molecule 1 |
| Socs3 | -0.48 | 5.18e-03 | -0.94 | 6.73e-10 | suppressor of cytokine signaling 3 |
| Trib3 | -0.96 | 4.72e-12 | 0.90 | 9.57e-12 | tribbles pseudokinase 3 |
| Autophagy |  |  |  |  |  |
| Hspa8 (Hsp70) | 0.53 | 5.09e-08 | -0.32 | 5.52e-05 | heat shock protein 8 |
| Map1lc3a (Lc3a) | 0.40 | 6.99e-04 | 1.02 | 1.97e-31 | microtubule-associated protein 1 light chain 3 alpha |
| Ulk1 | 0.41 | 3.77e-03 | -0.06 | 6.18e-01 | unc-51 like kinase 1 |
| Ulk2 | 0.26 | 3.97e-03 | -0.10 | 2.56e-01 | unc-51 like kinase 2 |

## DISCUSSION

In our study, we sought to identify novel glycans that might serve as optimal substrates for propionate production by gut microbes and enrich the gut microbiota in propiogenic bacterial taxa, to leverage the therapeutic potential of gut-derived propionate against CMRFs such as obesity, atherogenic dyslipidemia and insulin resistance. To do so, we screened a large library of synthetic glycans for propionate production using an e*x vivo* fermentation platform, and selected KB39 as glycan capable of supporting high propionate production. We showed that KB39 greatly increased the SCFAs acetate and propionate and increased the ratio of propionate over acetate and butyrate, both *ex vivo*, in fecal communities from healthy subjects and overweight patients with T2DM and *in vivo*, in western diet-fed *Ldlr^-/-^* mice. In the latter, KB39 demonstrated multiple beneficial effects, including a large decrease in blood total cholesterol and LDL-cholesterol, and a strong reduction in hepatic cholesterol and triglycerides, with only a mild attenuation in body weight gain. Those effects translated to a significant reduction in liver steatosis and inflammation, and a marked reduction in atherosclerosis, which were similar or superior to fenofibrate. In addition to its effect on lipids, KB39 strongly decreased fasting blood insulin, suggesting that KB39 increased insulin sensitivity. This is of particular importance considering the crucial role played by insulin resistance in the onset of T2DM and NAFLD and the increased risk of CVD ^66^.

To our knowledge, KB39 is the first synthetic glycan specifically selected for high propionate production by the gut microbiota and offers significant advantages *versus* oral propionate. Unlike propionate that has poor taste and smell and is readily absorbed in the upper gastrointestinal tract, KB39 is tasteless and odorless, non-absorbable, non-digestible and will be metabolized in the lower small intestine and colon, the natural sites of propionate absorption and signaling through its molecular receptors. In addition, and unlike oral propionate, KB39 has the potential to elicit beneficial changes in the composition of the gut microbiome of patients. KB39 enriched propiogenic bacterial taxa belonging to the *Bacteroidetes* phylum, including *Bacteroides, Parabacteroides* and *Alistipes*. Consistent with *Bacteroidetes* using the succinate pathway to generate propionate from complex polysaccharides, KB39 treatment also led to an increase in total and relative cecal succinate in the *Ldlr^-/-^*mice *in vivo*. Succinate itself is emerging as another important microbial metabolite improving metabolic dysfunctions ^67, 68^. In addition to *Bacteroidetes*, KB39 also increased the relative abundance of *Akkermansia muciniphila in vivo*, a microbe which has been shown to improve metabolic disorders and obesity through the production of immunomodulatory proteins (such as Amuc_1100) and improve the gastrointestinal barrier function ^69, 70^. KB39 also decreased bacterial species considered pro-inflammatory such as *Escherichia coli, Enterococcus faecalis* and *Klebsiella pneumoniae*. This, along with the beneficial effects of propionate, could play a role in KB39 anti-inflammatory properties observed in the liver. One potential mechanism of action for KB39 on lipid and glucose metabolism is the activation of the AMP-activated protein kinase (AMPK) as it is now established that AMPK is phosphorylated in response to propionate and other SCFAs in muscle, liver and intestinal tissue and AMPK is emerging as an important component in SCFA-conferred metabolic benefits ^20, 71, 72^.

In conclusion, we have shown that it is possible to identify synthetic glycans that support high propionate production through *ex vivo* screening and that administration of such glycans to western diet-fed *Ldlr^-/-^* mice results in a similar induction of high propionate producing gut microbiota *in vivo*. This has multiple beneficial effects, in lowering circulating LDL-cholesterol, reduction in fat accumulation in the liver and reduction in development of atherosclerotic lesions in multiple vessels throughout the body. These data highlight the potential of propionate producing glycans such as KB39 in improving major CMRFs and the prevention of cardiometabolic diseases.

## ACKNOWLEDGMENTS

We thank the Kaleido Biosciences’ Discovery and Technical Operations teams for their hard work and expertise and Peter Turnbaugh (UCSF) for critical reading of the manuscript.

## SOURCES OF FUNDING

This study was funded by Kaleido Biosciences.

## DISCLOSURES

C. Ronald Kahn is on the Kaleido Biosciences Scientific Advisory Board. All other authors either are or were employees of Kaleido Biosciences.

## SUPPLEMENTAL MATERIAL

Figures S1-S7

