## Supplementary material for "Modulation of the Gut Microbiome by Novel Synthetic Glycans for the Production of Propionate and the Reduction of Cardiometabolic Risk Factors"

### Figure S1

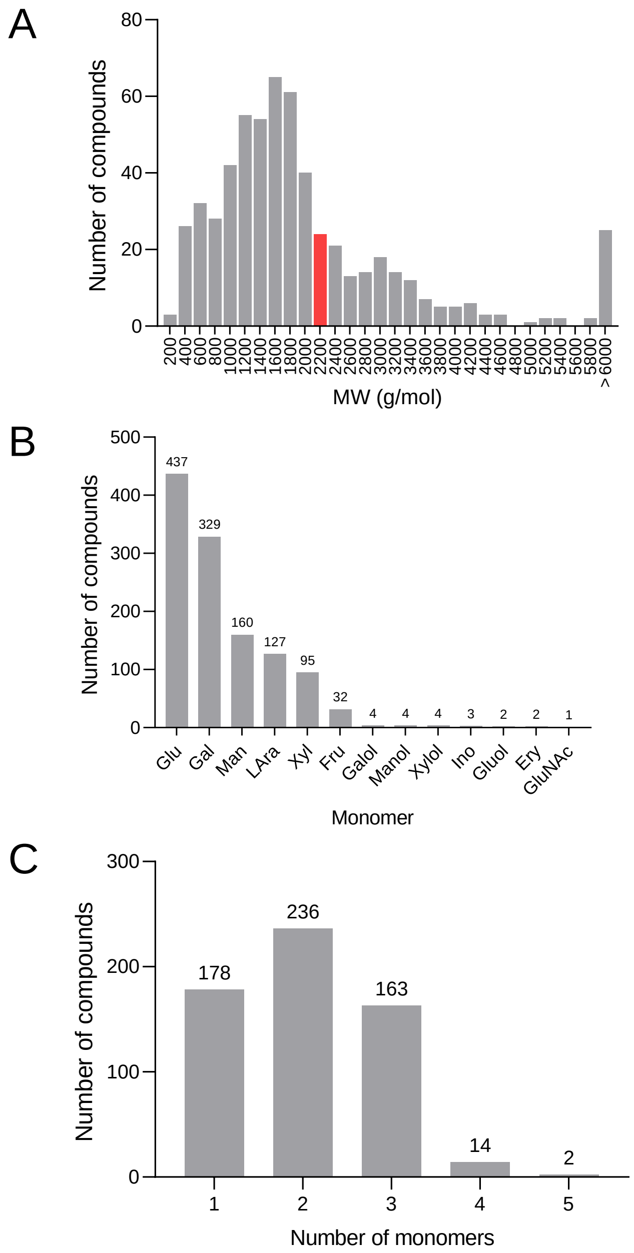


**Figure S1. Chemical features of the screened glycan library. (A)** Distribution of the molecular weight (MW) of tested glycans. The red bar indicates where KB39 is located. **(B)** Sugar monomers present in the chemical composition of the tested glycans. **(C)** Number of unique sugar monomers present in the chemical composition of the tested glycans. Glu: D-glucose; Gal: D-galactose; Man: D-mannose; Lara: L-arabinose; Xyl: D-xylose; Fru: D-fructose; Galol: galactitol; Manol: mannitol; Xylol: xylitol; Ino: inositol; Gluol: glucitol; Ery: erythritol; GluNAc: N-acetylglucosamine.

### Figure S2

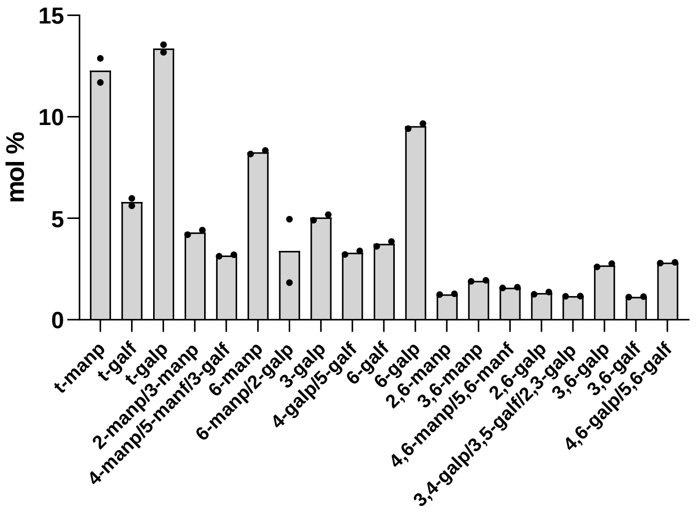


**Figure S2. Linkage analysis of KB39.** Mole percent of different glycosidic linkages in KB39. Labels show all linkage types with greater than 1% mole percent. Each linkage was analyzed in duplicate. man: mannose; gal: galactose; t: terminal; p: pyranose; f: furanose.

### Figure S3

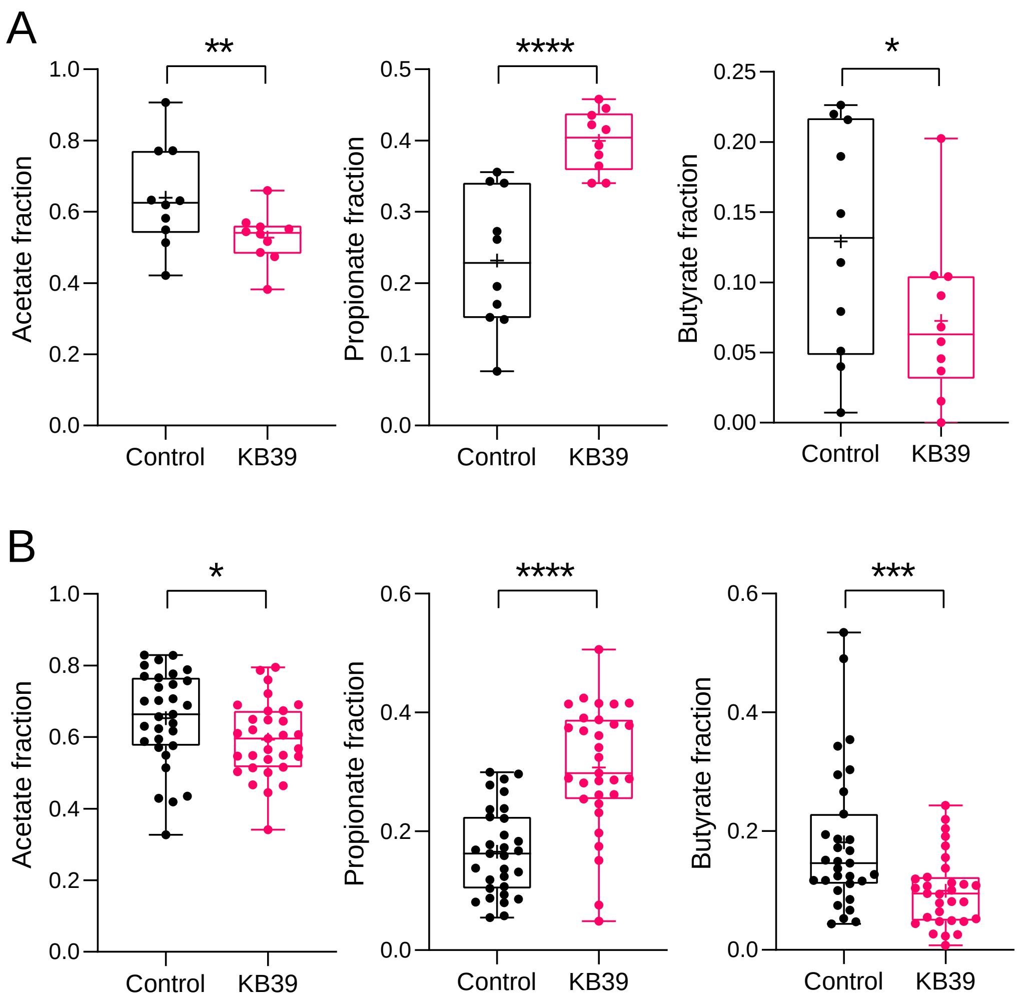


**Figure S3. Acetate, propionate and butyrate fraction of total SCFAs in human fecal microbiota treated with KB39 *ex vivo***. Fecal microbiota from 10 healthy subjects **(A)** and 31 overweight subjects with T2DM **(B)**. Fecal microbiota cultures were incubated without (negative control) or with KB39 and SCFAs were measured in culture supernatants. * $P<0.05$, ** $P<0.01$, *** $P<0.001$, **** $P<0.0001$, paired *t*-test.

### Figure S4

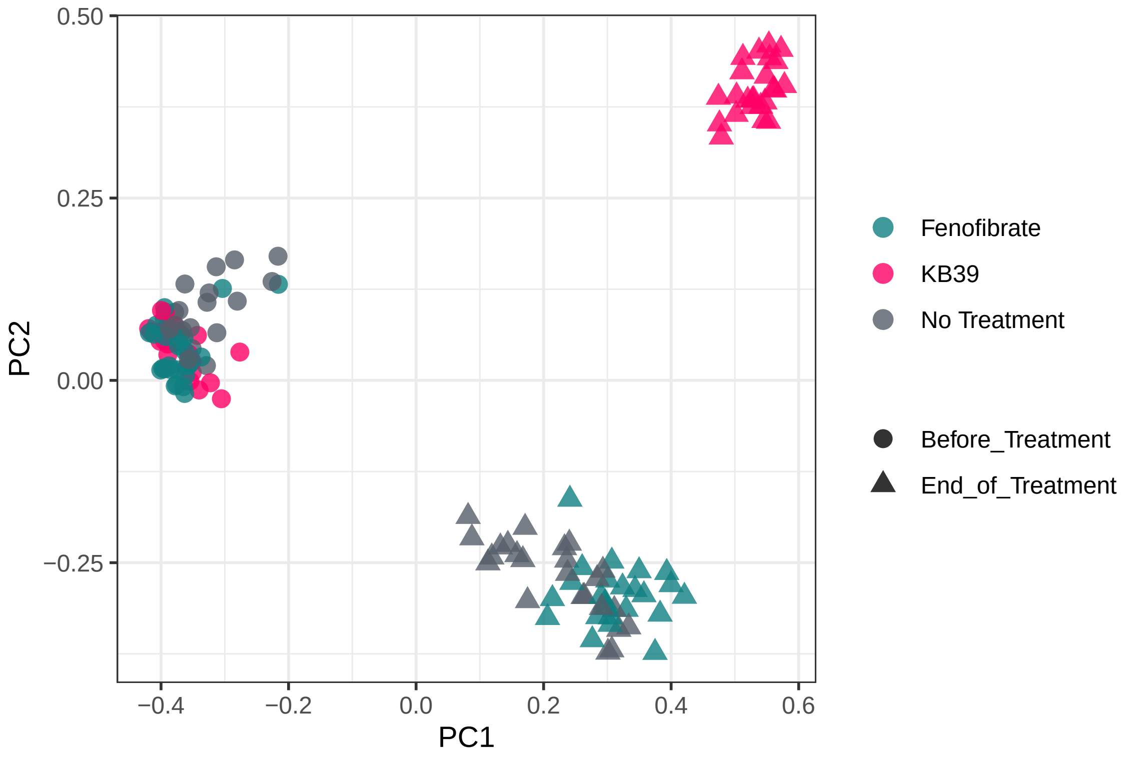


**Figure S4. Principal coordinate analysis on microbiome dissimilarity in non-treated, fenofibrate or KB39-treated western diet-fed *Ldlr*$^{-/-}$ mice.** Male *Ldlr*$^{-/-}$ mice were fed a western diet, western diet supplemented with KB39 (7.5% w/w) or fenofibrate (100 mg/kg/day) for 16 weeks. Fecal samples were collected before treatment during the last week of the treatment period.

### Figure S5

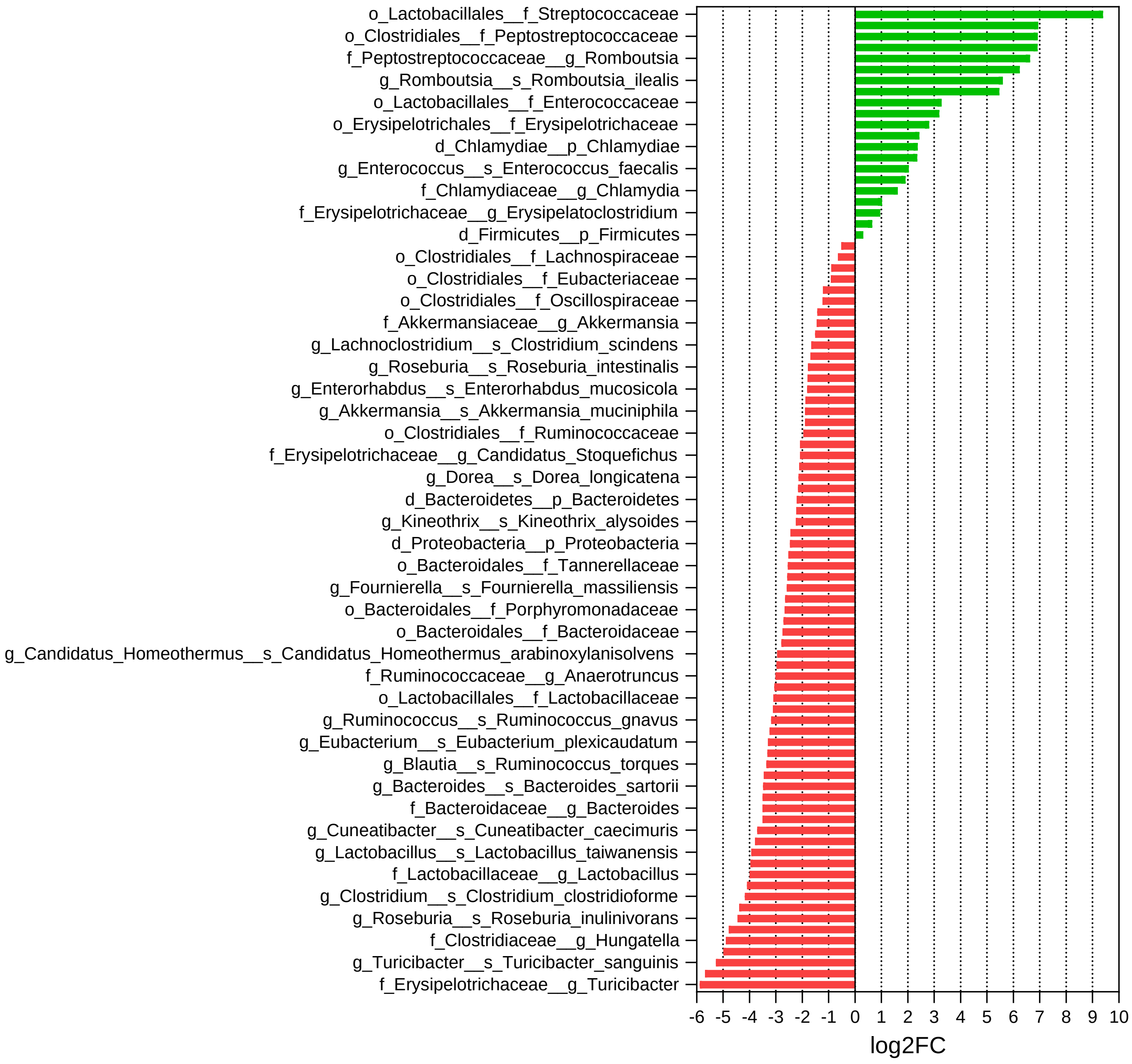


**Figure S5. Bacterial significantly enriched or depleted by western diet feeding in the gut microbiome of *Ldlr*$^{-/-}$ mice.** Male *Ldlr*$^{-/-}$ mice were fed a western diet for 16 weeks. Fecal samples were collected before switching the diet and during the last week of the treatment period. Significantly enriched and depleted taxa (fdr < 0.1, Wilcox rank sum test with FDR correction) at phylum, family, genus, and species levels between pre and post western diet feeding.

### Figure S6

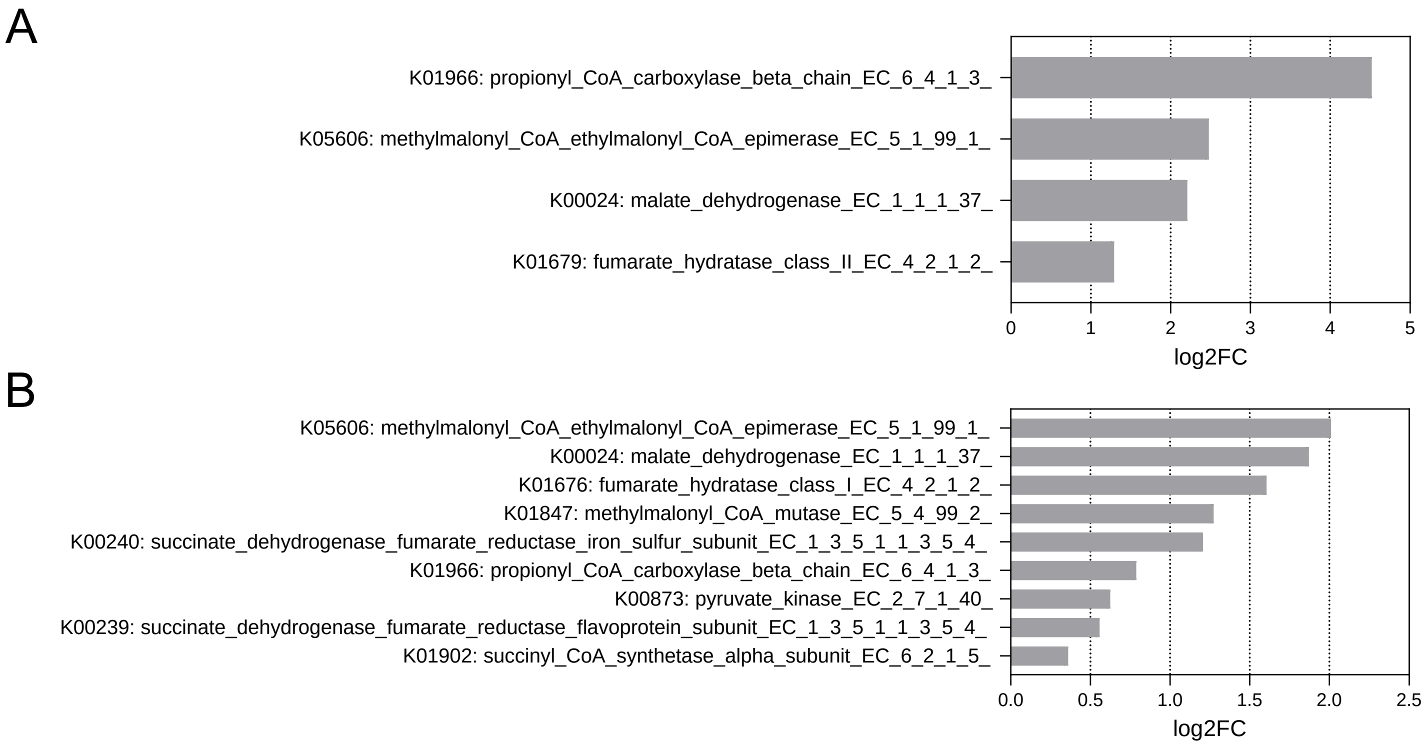


**Figure S6. Change in relative abundance for KO terms associated with propionate fermentation pathways *ex vivo* and *in vivo*. (A)** Fecal microbiota cultures from 31 overweight-T2DM subjects were incubated *ex vivo* without (negative control) or with KB39. Data represent the log2 fold change in relative abundance for KO terms associated with propionate fermentation pathways between the KB39 treated group and water (fdr < 0.1, mean log2 fold change > 0, Wilcox rank sum test with FDR correction). **(B)** Male *Ldlr*$^{-/-}$ mice were fed a western diet, or a western diet supplemented with KB39 (7.5% w/w) for 16 weeks. Fecal samples were collected during the last week of the treatment period. Data represent the log2 fold change in relative abundance for KO terms associated with propionate fermentation pathways between the KB39 and no treatment groups (fdr < 0.1, mean log2 fold change > 0, Wilcox rank sum test with FDR correction).

### Figure S7

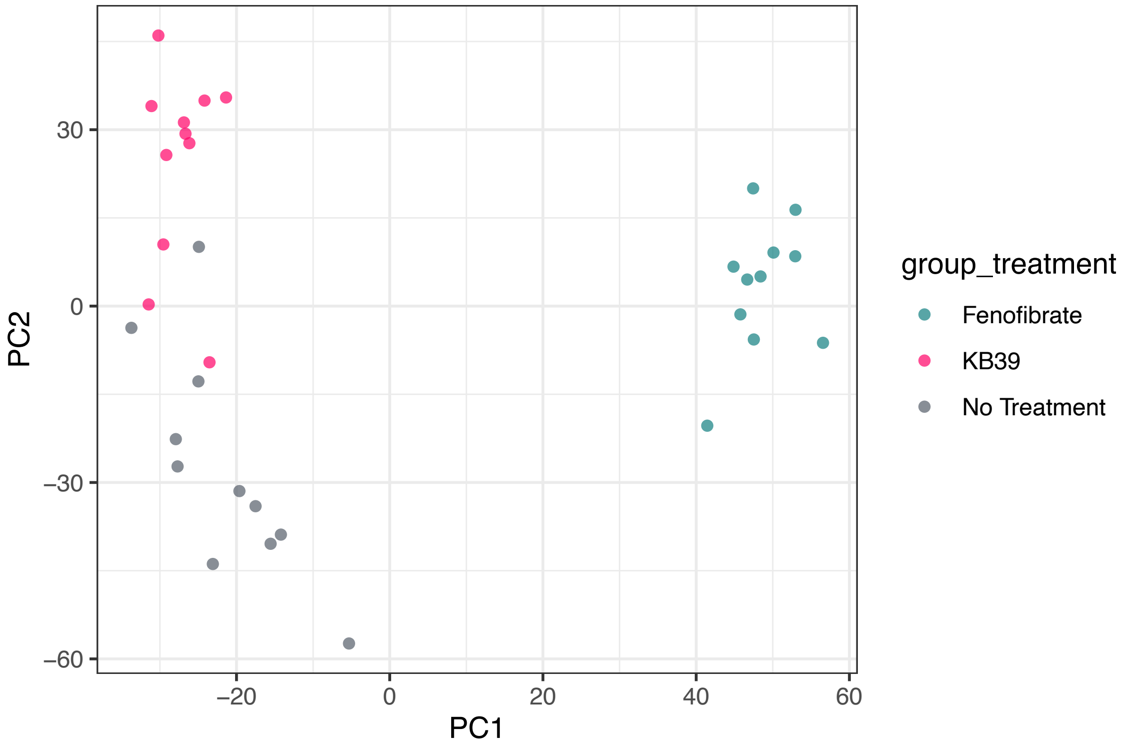


**Figure S7. Principal component analysis of liver gene expression profiles from western diet-fed *Ldlr$^{-/-}$* mice treated with KB39 or fenofibrate.** Male *Ldlr$^{-/-}$* mice were fed a western diet, western diet supplemented with KB39 (7.5% w/w) or fenofibrate (100 mg/kg/day) for 16 weeks. Livers were collected at termination and hepatic gene expression was analyzed by RNAseq. A principal component analysis (PCA) was performed on log transformed count per million values (logCPM).
